# Cholinergic modulation of rearing in rats performing a spatial memory task

**DOI:** 10.1101/2023.10.14.559618

**Authors:** Skylar Cassity, Irene Jungyeon Choi, Billy Howard Gregory, Adeleke Malik Igbasanmi, Sarah Cristi Bickford, Kiara Tyanni Moore, Anna Elisabeth Seraiah, Dylan Layfield, Ehren Lee Newman

## Abstract

Spatial memory encoding depends in part on cholinergic modulation. How acetylcholine supports spatial memory encoding is not well understood. Prior studies indicate that acetylcholine release is correlated with exploration, including epochs of rearing onto hind legs. Here, to test whether elevated cholinergic tone increases the probability of rearing, we tracked rearing frequency and duration while optogenetically modulating the activity of choline acetyltransferase containing (i.e., acetylcholine producing) neurons of the medial septum in rats performing a spatial working memory task (n = 17 rats). The cholinergic neurons were optogenetically inhibited using halorhodopsin for the duration that rats occupied two of the four open arms during the study phase of an 8-arm radial arm maze win-shift task. Comparing rats’ behavior in the two arm types showed that rearing frequency was not changed but the average duration of rearing epochs became significantly longer. This effect on rearing was observed during optogenetic inhibition but not during sham inhibition or in rats that received infusions of a fluorescent reporter virus (i.e., without halorhodopsin; n = 6 rats). Optogenetic inhibition of cholinergic neurons during the pre-trial waiting phase had no significant effect on rearing, indicating a context-specificity of the observed effects. These results are significant in that they indicate that cholinergic neuron activity in the medial septum is correlated with rearing not because it motivates an exploratory state but because it contributes to the processing of information acquired while rearing.

## Introduction

Determining the mechanisms that support the encoding of new memories remains a major outstanding challenge. Acetylcholine is important for memory encoding (Whishaw et al., 1985; Blokland et al., 1992; Elvander et al., 2004; Martyn et al., 2012; but also see Parent & Baxter, 2004) and its release is well correlated to exploratory behaviors including rearing up onto hind legs in rodents (Acquas et al., 1996; Thiel et al., 1998; Giovannini et al., 2001; Kopsick etl a., 2022). Yet, it remains poorly understood what role acetylcholine has in regulating exploration. Does high cholinergic tone motivate exploration to drive information acquisition? Or, alternatively, is exploration motivated by a different mechanism and acetylcholine plays a downstream role supporting the encoding of the newly acquired information? This project sought to test between these possibilities.

A dominant hypothesis for how acetylcholine supports memory encoding is that it enacts a switch of functional modes, promoting processing of bottom-up, sensory, information (for relevant reviews, see Hasselmo, 2006; Newman et al., 2012; Honey, Newman, Schapiro, 2017). This hypothesis is supported by circuit physiology studies indicating that cholinergic modulation increases the relative strength of synapses from bottom-up projections relative to lateral projections (e.g., Hasselmo & Schnell, 1994; Giocomo et al., 2005; Newman et al., 2013; Douchamps et al., 2013; Dannenberg et al., 2015). This switch in the circuit dynamics allows high-fidelity encoding of information carried into the circuit by extrinsic fibers by limiting interference from intrinsic projections. What is not addressed by this hypothesis is whether acetylcholine, beyond modulating the circuit dynamics, is also responsible for aligned exploratory behaviors.

The role of acetylcholine in modulating exploratory behaviors is not well understood. Existing data indicate that cholinergic tone and exploratory behavior are functionally related. Supporting evidence comes from the study of cholinergic tone during the initial exploration of novel enclosures (Acquas et al., 1996; Thiel et al., 1998; Giovannini et al., 2001) and activity of acetylcholine producing neurons of the medial septum and diagonal band in freely exploring mice (Kopsick et al., 2022). These studies reveal a consistent and robust positive relationship between exploration and cholinergic signaling. Pharmacological modulation studies involving local infusions of the cholinergic agonist carbachol into the septum provide mixed results (Monmaur et al., 1997; cf. Elvander et al., 2004). Monmaur et al. (1997) observed that, following agonist infusion, rats reared onto their hind legs more frequently when left in a narrow open cylinder without an overt task. Elvander et al. (2004) did not replicate this effect. However, they used a larger testing enclosure and instead found a significant increase in the total distance traveled which may, nonetheless, reflect increased exploration. Neither study examined exploration in the context of an overt memory paradigm leaving it unknown what relationship these changes had on memory encoding. Because rearing can be motivated by different objectives, for example, to plot an escape or acquire information regarding distal cues and environmental boundaries (Lever et al., 2006), to understand the relevance for spatial memory encoding, it is important to test the relationship between cholinergic modulation and exploration in the context of a spatial memory task. Rearing during the study phase of the 8-arm radial maze win-shift task is a relevant key epoch of spatial memory encoding (Layfield et al., 2023) making this a task well suited to the goals of the present study.

We hypothesized that the functional relationship between exploration and cholinergic tone could exist for two possible reasons. In the first, acetylcholine plays an upstream role, generating an exploratory behavioral state wherein locomotion and rearing become more likely. In the second, exploration is caused by other mechanisms that may also trigger acetylcholine release and the increased cholinergic tone supports the encoding of information acquired during the exploratory bout. Though not strictly mutually exclusive, we reasoned that these alternative hypotheses could be dissociated by examining exploratory behavior during optogenetic modulation of medial septal cholinergic neurons. If these neurons support the generation of an exploratory state, inhibition of their activity would decrease the rate and extent of exploratory behaviors. Alternatively, if these neurons support the encoding of acquired information rather than generate the exploratory state itself, inhibition of their activity could increase exploratory behavior in compensation for the impediment to the encoding process.

To test between these hypotheses, we used CRE-dependent viral transfection of halorhodopsin in choline acetyltransferase expressing neurons (ChAT neurons) of the medial septum to modulate the cholinergic tone in downstream targets. Cholinergic neurons of the medial septum (MS) were targeted as these are the major projection of acetylcholine to the hippocampus (Leranth & Frotscher, 1985), which is the putative site of spatial memory encoding in the 8-arm win-shift paradigm (Olton & Samuelson, 1976; but also see Layfield et al., 2020). In the 8-arm radial maze win-shift task, rats collect sugar pellets from four random arms in the study phase and, after being removed from the maze briefly, are given access to all eight arms in a test phase wherein sugar pellets can be found on the four arms not previously visited. In the study phase only, the activity of the ChAT neurons was inhibited through light stimulation throughout the time rats visited two of the four arms. Exploratory behaviors were compared between these ‘real stimulation arms’ and the remaining two ‘no-stimulation arms.’ The use of optogenetic techniques permitted within-trial manipulations, allowing us to compare rearing in real-stimulation and no-stimulation arms to test for rapid modulation of exploratory behavior.

Additionally, a between-trial but within-subject manipulation wherein the laser was either powered on (real-stimulation) on half of the trials and the laser remained off (sham-stimulation) for the other half allowed us to compare the influence of optogenetic inhibition on task performance. Finally, a between-subject manipulation of virus type transfected rats with either halorhodopsin to implement the inhibition or fluorescent reporter only to control for possible confounds. We show that optogenetic inhibition of medial septal cholinergic neurons did not decrease any exploratory behavior and, instead, increased rearing duration. These results are consistent with the hypothesis that cholinergic tone supports the encoding of newly acquired information but does not generate an exploratory state.

## Methods

All procedures and surgeries were conducted in strict accordance with National Institutes of Health and under a protocol approved by the Indiana University Institutional Animal Care and Use Committee (IU IACUC).

### Subjects

A total of 27 *ChAT*:: *Cre*+ Long Evans rats (Witten et al., 2011) were used for this study. *ChAT::Cre+* transgenic rats express Cre recombinase in choline acetyltransferase (ChAT) positive neurons allowing for cell-type specific expression of optogenetic tools (Witten et al., 2011). Here, *ChAT::Cre+* rats were used for the selective expression of halorhodopsin in cholinergic neurons in the medial septum. Rats were bred in house and identified as CHAT::Cre+ by ear punch genotyping (TransnetYX). The total number of 27 rats consisted of 17 rats transfected with halorhodopsin, six that were transfected with a fluorescent reporter only, three that failed to reach behavioral criterion, and one that failed to thrive following surgery. Of the 17 rats that were transfected with halorhodopsin, nine (5 male) received optical cannula implants while eight (3 male) were implanted with optical cannula and microelectrode arrays. Of the six transfected with fluorescent reporter, four (2 male) were implanted with optical cannula only and two (1 male) were implanted with optical cannula and microelectrode array. The data obtained from the microelectrodes did not contribute to this project. Rats were individually housed with ad libitum access to water and food restricted to ∼90% of free-feeding weight. One female was removed from the study for failing to reach behavioral criterion. All were maintained on a 12-hour light / dark cycle.

### Apparatus

Behavioral training took place in a custom 8-arm radial maze with computer-controlled pneumatic drop doors at the entrances. The maze had a 33.2 cm wide hub and each arm measured 48.26 cm long, 10.79 cm wide with 20.95 cm tall walls. Walls were made from clear acrylic to allow for visual orientation to distal cues that surrounded the maze. The floors were made from opaque matte white acrylic. The maze was open-topped to allow testing of rats tethered by the fiber optic patch cable. At the end of each arm were food wells in which 45 mg sucrose pellets (Bio-Serv, Flemington, NJ) were delivered. The maze was situated in a 3m by 3.6m room, surrounded by rich visual cues to facilitate allocentric orientation. Cues were constructed from brightly colored tape or construction paper with a row of 6 cues evenly spaced level with the walls of the maze, and a second row of cues approximately 2 feet above the first row of cues. The room also contains a framed image, a desk, a bookshelf, and the holding pedestal on which rats were placed between the trial phases immediately adjacent to the maze (see photo in Fig 1). The experimenter was present in the room during all sessions, standing at the same location and wore the same PPE (white lab coat) during all sessions to maintain a stable distal cue. A high-definition camera (FLIR Flea3 or Intel RealSense D435) mounted to the ceiling above the maze were connected to a PC running Open Broadcast Studio (OBS) to record the movements of the rat for offline analysis. A CE:YAG laser diode optical head laser system (Doric Lenses) filtered to 570 nm to 615 nm was mounted to the ceiling. A 400 μm core patch cord connected the laser head to the animal through a passive optical rotary to allow for free animal rotation. Surplus cord was restrained by elastic strings to enable free movement of the animal while keeping the cord out of the way.

**Figure 1:**
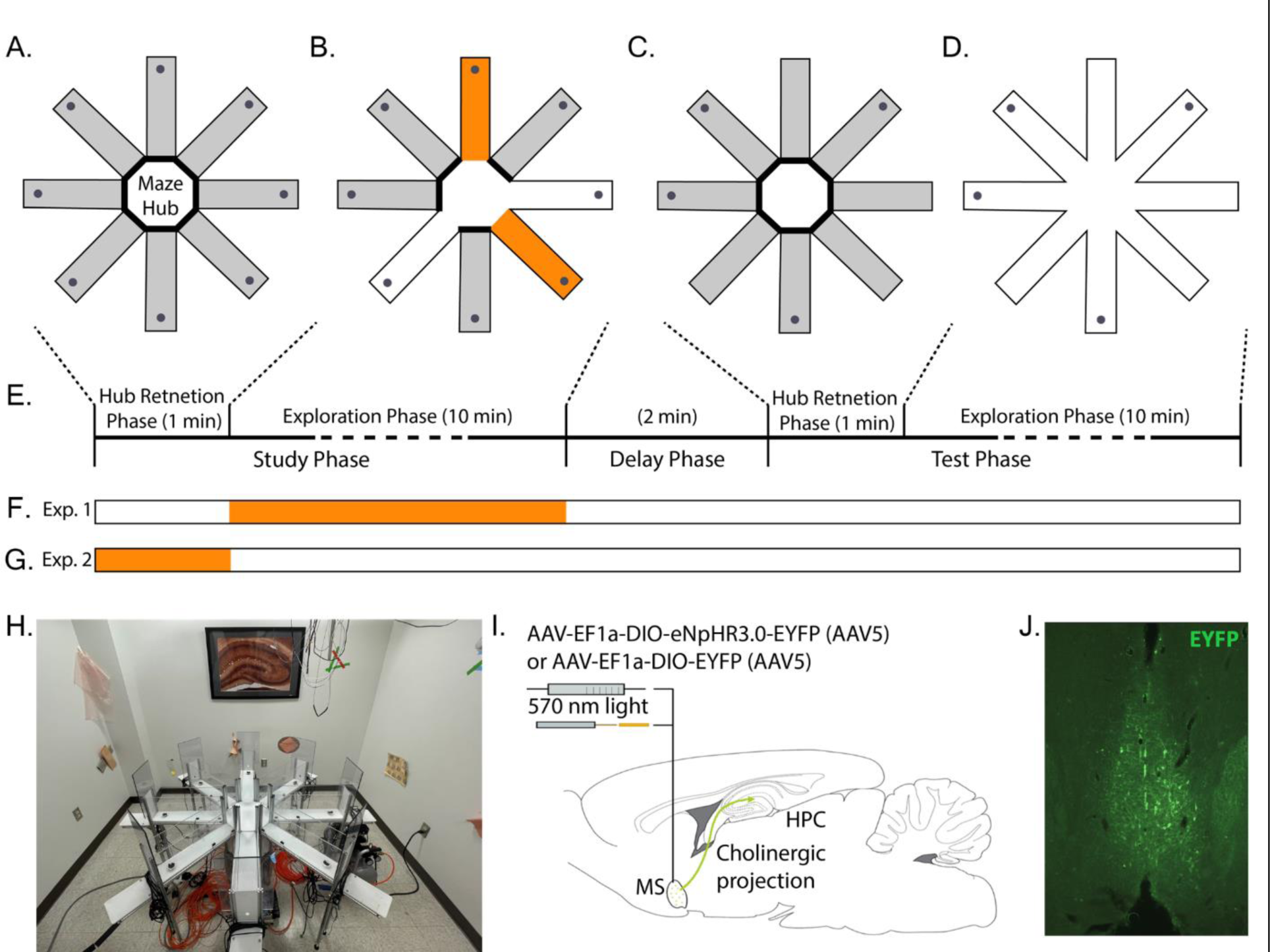
Overview of experimental approach. A-E. Rats performed a win-shift task on the 8-arm radial arm maze. A. Trials began with a *pre-trial hub retention phase* wherein the rat was placed in the hub with doors to all arms closed for 1 min. B. *Study phase*. A different set of four random doors opened on each trial. Rats could collect sucrose pellets from each (gray dots at arm ends). Two arms were randomly selected to be ‘stim-arms’ (orange) and the remaining two were ‘non-stim-arms’ (white). The laser was triggered throughout the time rats spent in in stim-arms. Rats were removed from the maze after collecting all pellets. C. After a 2 min. delay, rats were replaced in the hub with all doors closed for another 1 min. retention phase. D. A *test phase* began when all eight doors opened. Rats could collect pellets from arms not visited during the study phase. E. Timeline of a single trial. The times indicated for each exploration phase indicate the maximum allowed. Each phase was terminated once all available pellets were collected. F. Indication of timing of possible light delivery during real-stim trials. G. Indication of timing of light delivery during hub-stim trials. H. Photograph of experimental apparatus showing availability of distal cues around maze. I. Illustration of optogenetic modulation approach. ChAT::CRE+ rats were infused with either a virus packaged with CRE-dependent halorhodopsin and fluorescent reporter or a virus packaged with CRE-dependent reporter. The infusion targeted the medial septum, the dominant source of cholinergic input to the hippocampal formation (green arrow). An optical fiber was implanted just dorsal to the medial septum. J. Representative example of the distribution of EYFP expression in a rat from this study infused with CRE-dependent halorhodopsin and reporter.

### Behavioral training

#### Habituation & Preliminary Training

Rats were handled for ten minutes and given 20–30 sucrose pellets to habituate them to experimenter handing and rewards for 3 days prior to preliminary training. Preliminary training lasted for three days, with one session per day to introduce collection of sugar pellet rewards from the 8-arm maze. Three sugar pellets were placed along all eight maze arms leading from the maze hub to feed dishes baited with two sugar pellets. Rats were placed in the maze hub of the maze with all doors open and allowed to forage for ten minutes or until all sugar pellets were consumed. After each session, maze floors, walls, and doors were cleaned with 10% chlorohexidine solution to ensure cleanliness of the maze and reduce odors that might provide smell cues.

#### Initial training

Rats underwent initial training for ten days, one session/day and one trial/session. During each, rats were trained that all arms were baited with two sugar pellets each day. Training began when the rat was placed in the maze hub and every door was promptly opened. Rats were allowed to forage until all pellets were collected or 15 minutes had elapsed, whichever came first. The time the rat was kept in the maze hub before each door opened was increased incrementally in following sessions by ten seconds until one minute was reached.

#### Task

The spatial memory task used here was the delayed-win-shift task on an 8-arm radial arm maze. The task consisted of three phases, a study phase, a delay phase, and a test phase as shown in Fig. 1A. Prior to the study phase, the rat was placed in the maze hub with all doors shut for 20-60 seconds. After the delay, a random set of four doors opened and the rat was allowed to collect pellets from each. The rat was then removed from the maze and placed on a pedestal next to the maze for a two-minute delay phase. During the delay, the maze was cleaned with chlorhexidine. The test phase began by placing the rat in the maze hub with all doors shut for 60 seconds. After 60 seconds, all eight doors opened. The four arms that had not opened during the study phase baited food wells. The test phase ended after all the pellets had been consumed or when 15 min elapsed. Regular training (∼5 days/week) continued until rats achieved behavioral the criterion of no more than three errors over four days.

### Behavioral scoring

The primary dependent measures of spatial memory performance were percent correct and number of arm entries. Percent correct was measured as the number of the first four arm entries in the test phase that contained rewards divided by four. Number of arm entries was measured as the total number of arms entered to find all four baited reward sites in the test phase. A visit to an arm was defined to occur when the rat’s hind feet entered the arm. An arm visit counted as an error if the arm had been visited earlier in the trial. Secondary analyses also examined ‘reentries.’ Reentries were quantified as the number of times in the test phase that a rat entered an arm that had been open in the study phase arm.

### Exploration scoring

The rat position was tracked by applying DeepLabCut (Mathis et al., 2019; Nath et al., 2019) to video recordings of the rat in the maze collected from an overhead camera (RGB sensor of a RealSense Depth Camera D435; Intel). The camera captured the whole of the maze at 30 Hz and 640 x 480 resolution. A DeepLabCut network was built with 24 frames from each of the 10 rats (240 total frames) labeled to mark the nose, left ear, right ear, and tail base (rump). Examples are shown in Fig. 1B. After an initial ∼300k training iterations, an additional 240 ‘jumpy’ frames were extracted, relabeled, and added to the training set. The network then received another ∼600k training iterations on the expanded training set. Network performance was then confirmed by manual inspection by checking labeled videos and quantifying jumpy frames in the extracted tracking data. Prior to use, the tracking data was preprocessed to interpolate over jumps of >5 cm from one frame to the next. Rat position (e.g., summing up total movement) was estimated by the position of the rump. Rearing events were manually scored through offline analysis by experimenters. Rearing events began once at least one forepaw left the maze floor and ended once both forepaws returned to the ground. Experimenters were blinded to the experimental hypothesis but could not be blinded to when optogenetic stimulation was applied as the laser light could be seen in the videos.

### Surgery

After meeting behavioral criterion, rats underwent stereotaxic surgery to allow optogenetic inactivation of medial septal cholinergic neurons. Pain and discomfort were minimized via administration of buprenorphine (0.03mg/kg, SC) and meloxicam (2mg/kg, SC) immediately before surgery onset and a surgical plane of anesthesia was maintained with isoflurane (1.25-2.5%) following induction (4%). Once induced, the scalp of the rat was shaved and it was placed into a stereotaxic frame (Kopf Instruments). A scalp incision from anterior to posterior was made across the skull. A craniotomy for the viral injection and optical fiber was placed at [AP +06. mm, ML +1.0 mm]. A 10 μl gastight syringe with 32 ga needle (Hamilton) connected to a microinjection robotic stereotaxic arm (Neurostar) at a 9° angle laterally in the sagittal plane to avoid midline vascular structures was used to perform a microinjection of 2.5 μl to the medial septum [AP +0.6mm, ML ∼0mm, DV 6.1mm]. Individual rats received either a virus carrying halorhodopsin and fluorescent reporter (AAV(5)-EF1a-DIO-eNpHR3.0-EYFP) (UNC Vector Core) or fluorescent reporter only (AAV(5)-EF1a-DIO-EYFP) (UNC Vector Core). An optical cannula (NA=0.66) was lowered into the same area as the needle, delivered to rest 300-500 microns above the viral injection and fixed with bone screws inserted into the skull with dental acrylic. The site was suture closed around the implant. Rats recovered for 7 days before moving on to the next phase. A summary is shown in Fig. 1C.

### Optogenetic control

Optogenetic control was implemented with the eNpHR3.0 halorhodopsin which inhibits neural activity with photostimulation (Gradinaru et al., 2010). Activation of halorhodopsin was achieved using light from a CE:YAG laser diode optical head laser system (Doric) filtered to 570 - 615 nm. Laser output was delivered to the medial septum by way of a patch fiber connected to a rotary joint (Doric) and, from the rotary joint a dual fiber optic patch cord (Doric) that was coupled to the implanted optical fibers prior to each testing session (core diameter = 400 um; numerical aperture = 0.57). Light intensity was controlled by the Doric Neuroscience studio software to obtain 5-10 mW at the tip of the fiber in the brain using a photodiode power sensor coupled to a power meter (Thorlabs). Laser activation was triggered by the experimenter via a trigger button controlled TTL signal generated by an Arduino unit (Arduino due). Rats were connected to the optical patch cords throughout all phases of testing regardless of experimental condition.

### Experimental conditions

Both within trial and between trial experimental manipulations were used. The within-trial manipulation consisted of ‘stim-arms’ and ‘no-stim-arms.’ Stim-arms were two randomly selected arms (using Random.org), of the four arms opened in the study phase, wherein the experimenter pressed a button to trigger the laser. The button remained pressed for the duration that the rat’s head remained in the designated arms. The button was not pressed when the rat was in the no-stim-arms. In no condition was optogenetic inhibition activated in any phase other than the study phase. The within-trial manipulation allowed for temporally precise analysis of whether laser activation changed acute exploratory behavior. The between trial conditions consisted of real-stim, sham-stim and hub-stim trials. In real-stim trials, the laser was powered on and delivered light when the trigger button was pressed. In sham-stim trials, the laser remained off and no light was delivered when the trigger button was pressed. In hub-stim trials, the laser was powered on and was triggered throughout the pre-trial hub retention phase (see Fig 1). The between-trial manipulation allowed for analysis of whether laser activation either during the task (real-stim vs. sham-stim) or prior to the task (hub vs. sham-stim) changed exploratory behavior or task performance across trials.

Two cohorts of rats were run through these conditions. The first, consisting of nine halorhodopsin rats with optical fibers, completed 10 of each of the real-stim and sham-stim trials. Six then completed 8-10 trials of each of the hub and sham-stim trials. The second cohort, consisting of eight halorhodopsin with optical fibers and microelectrode arrays and six reporter-only rats with optical fibers (half with microelectrode arrays), completed 3-5 of each of the trial types (real-stim, sham-stim and hub) in a randomized interleaved fashion.

### Histology

Upon completion of testing, animals were euthanized via isoflurane overdose and perfused intracardially with phosphate buffered saline (PBS) followed by a 4% paraformaldehyde saline solution. Brains were saturated with a 30% sucrose solution prior to sectioning. Coronal sections (40 um thick) were cut with a microtome (American Optic company). Immunohistochemistry was performed on free-floating sections to amplify induced enhanced yellow fluorescent protein (EYFP) reporter signaling. Sections were first rinsed with PBS then blocked with buffer (PBS, 5% normal goat serum, and 0.4% Trition X-100). This was followed by overnight incubation with conjugated anti-green fluorescent protein (anti-GFP) rabbit antibody (1:2000; catalog no. A21311; Invitrogen). Finally, the sections were rinsed with PBS, mounted on slides, and cover slipped with DAPI and Fluoroshield. Labeled sections were imaged with an epifluorescent microscope. An example section is shown in Fig. 1D.

### Statistical Analyses

Hypothesis testing was performed using generalized linear mixed effects (glme) modeling. This approach allowed for hierarchical modeling of the effects given the different numbers of trials across animals and conditions. Tests related to rearing (count, duration), exploration (distance traveled, dwell time), and performance (percent correct, arm entries, arm reentries) used the equation *’met ∼ opto + (1|rat)’* where *met* is a stand-in variable for each metric of interest (rear count, duration, etc.), *opto* labels trials as real-stim versus sham-stim, and *rat* labels data obtained from each rat. This equation implements a random intercept model across rats with a fixed effect slope for the stimulation type. Tests for interactions between rat cohort (halorhodopsin versus reporter-only) and trial type (real-stim versus sham-stim) for each metric used the equation *‘met ∼ opto + (opto | cnd) + (1 | rat)’* where *met*, *opto*, and *rat* are as before and *cnd* indicates which cohort a rat is from. This equation models each metric with a random slope for opto across cohorts and random intercept across rats. Statistical results are reported with the glme estimated intercept coefficients for each condition, the glme estimated difference (indicated with glme est.), the 95% confidence interval on the difference (indicated in square brackets), the *t*-statistic with the degrees of freedom it was calculated over in parenthesis, and the corresponding *p* value. Across analyses, information is provided to the analysis about all trials and random effects are calculated over trials based on rats. Accordingly, the degrees of freedom reported in the manuscript is based on the total number of trials. Significance was defined at the α= 0.05 level.

## Results

Our goal was to test the hypothesis that cholinergic neurons of the medial septum regulate rearing behavior in a spatial memory task. Two cohorts of ChAT-CRE rats were trained to perform a win-shift task on the 8-arm radial arm maze. In the first cohort, a CRE-dependent virus carrying the DNA for halorhodopsin and a fluorescent reporter was surgically infused into the medial septum. In the second cohort, the infused virus contained only the DNA for the reporter. The first cohort served as the main experimental group while the second served as a control to ensure observed effects were dependent upon the halorhodopsin. Both cohorts performed three trial types. In ‘Real stimulation trials,’ the laser was powered on and the rats received light stimulation in two of the four accessible maze arms during the study phase. No stimulation was delivered in the other two arms. In the ‘Sham stimulation trials,’ the laser remained powered down so that no light was delivered to the rats when they entered the stimulation arms. The sham stimulation trials established the expected behavior when no manipulation was performed. Finally, in ‘Hub stimulation trials,’ light stimulation was delivered in the epoch after a rat was placed in the hub of the maze but before the doors opened. The hub stimulation trials established whether the effects of stimulation on rearing were specific to the task context or were general across contexts. Analyses tested for stimulation effects on the number and duration of rears as well as effects on the distance traveled, dwell times, and task performance. Our working hypothesis was that inhibiting ChAT neurons in the medial septum would reduce the number and duration of rears. Instead, as reported here, we found no change in the number of rears and a significant increase in rear duration.

To test the hypothesis that cholinergic activity and rearing are well correlated because the exploratory behavioral states are induced by increased cholinergic tone, we first asked if rearing frequency was significantly reduced when ChAT neurons of the medial septum were optogenetically inhibited. Toward this end, we compared the number of rears in arms wherein the laser was activated (‘real-stim-arms’) and those wherein the laser not activated (‘no-stim-arms’) in rats transfected with halorhodopsin. We found that the number of rears in real-stim-arms (glme est. = 1.2 rear) and in no-stim-arms (glme est. = 1.2 rear) were not significantly different (glme est. = 0.03 [-0.17 0.24], *t*(261) = 0.32, *p* = 0.75; Fig. 2A & 2B). No differences were expected in the other conditions and none were observed. Explicitly, no differences were observed in sham stimulation trials (sham stim = 1.0; no stim = 1.0; glme est. = 0.03 [-0.11 0.12], *t*(310) = 0.05, *p* = 0.96; Fig. 2A & 2B) nor in either trial type in the rats transfected with reporter only (real stim = 1.0 rear; no stim = 0.97 rear; glme est. = -0.03 [-0.32 0.25], *t*(56) = -0.24, *p* = 0.81; sham stim = 0.85 rear; no stim = 0.9 rear; glme est. = 0.05 [-0.21 0.32], *t*(56) = 0.39, *p* = 0.69; Fig. 2C & 2D). Our failure to find a change in rearing frequency indicates that medial septal ChAT neuron activity does not generate an exploratory behavioral state that is marked by increased rearing.

**Figure 2:**
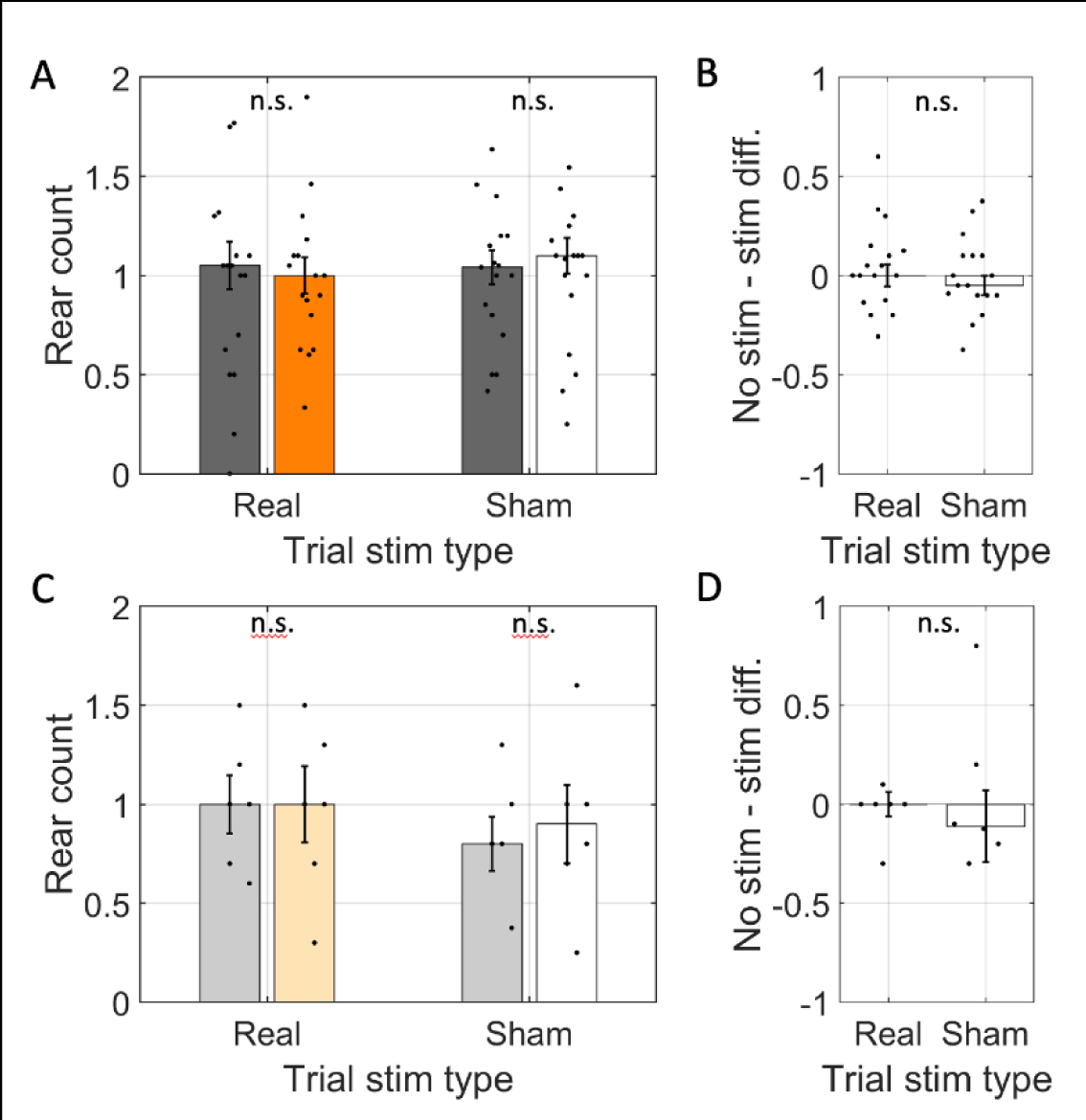
Rear frequency was not changed by optogenetic inhibition. A. The average number of rears per arm (rear count) across rats transfected with halorhodopsin (n=17). Rear count is shown separately for arms with simulation (orange / white) and arms without stimulation (gray) on trials with the laser powered on (Trial stim type: real) and trials with the laser powered off (Trial stim type: sham). B. The difference in average rear count between arms with and without stimulation in real and sham stimulation trials. Rear count was not different across arm types in either trial type and trial types were statistically indistinguishable. C & D. Same as A & B but for rats transfected with reporter only (n=6). In all panels, dots indicate the mean over trials for individual rats, bar height indicates the median over rats, and error bars indicate standard error over rats. N.s. indicates *p* > 0.05.

An alternative hypothesis to explain the correlation between rearing and cholinergic activity is that the processing that occurs during rearing is supported by acetylcholine. This hypothesis suggests that rears that occur in the absence of normal cholinergic activity would be altered. To assess this hypothesis, we asked if the mean duration of rearing events were affected by optogenetic inhibition. Indeed, we found that the mean rear duration in real-stim-arms (glme est. = 4.7 s) was significantly longer than in no-stim-arms (3.7 s; glme est. = 1.0 [0.31 1.7], *t*(241) = 2.83, *p* = 0.005; Fig. 3A & 3B). Note that the degrees of freedom of the test here is lower than that in other tests because trials with no relevant rears did not contribute to the calculation of the average rear duration.Sham stimulation did not significantly affect the duration of rears in the halorhodopsin rats (sham-stim = 4.4 s vs. no-stim = 4.0 s; glme est. = 0.35 [-0.31 1.01], *t*(294) = 1.05, *p =* 0.29; Fig. 3A & 3B). This was a significant difference between the real-stim and sham-stim trials (glme est. = 0.66 [0.11 1.20], *t*(534) = 2.4, *p* = 0.017), indicating that real stimulation lengthened rear duration relative to sham stimulation.

**Figure 3:**
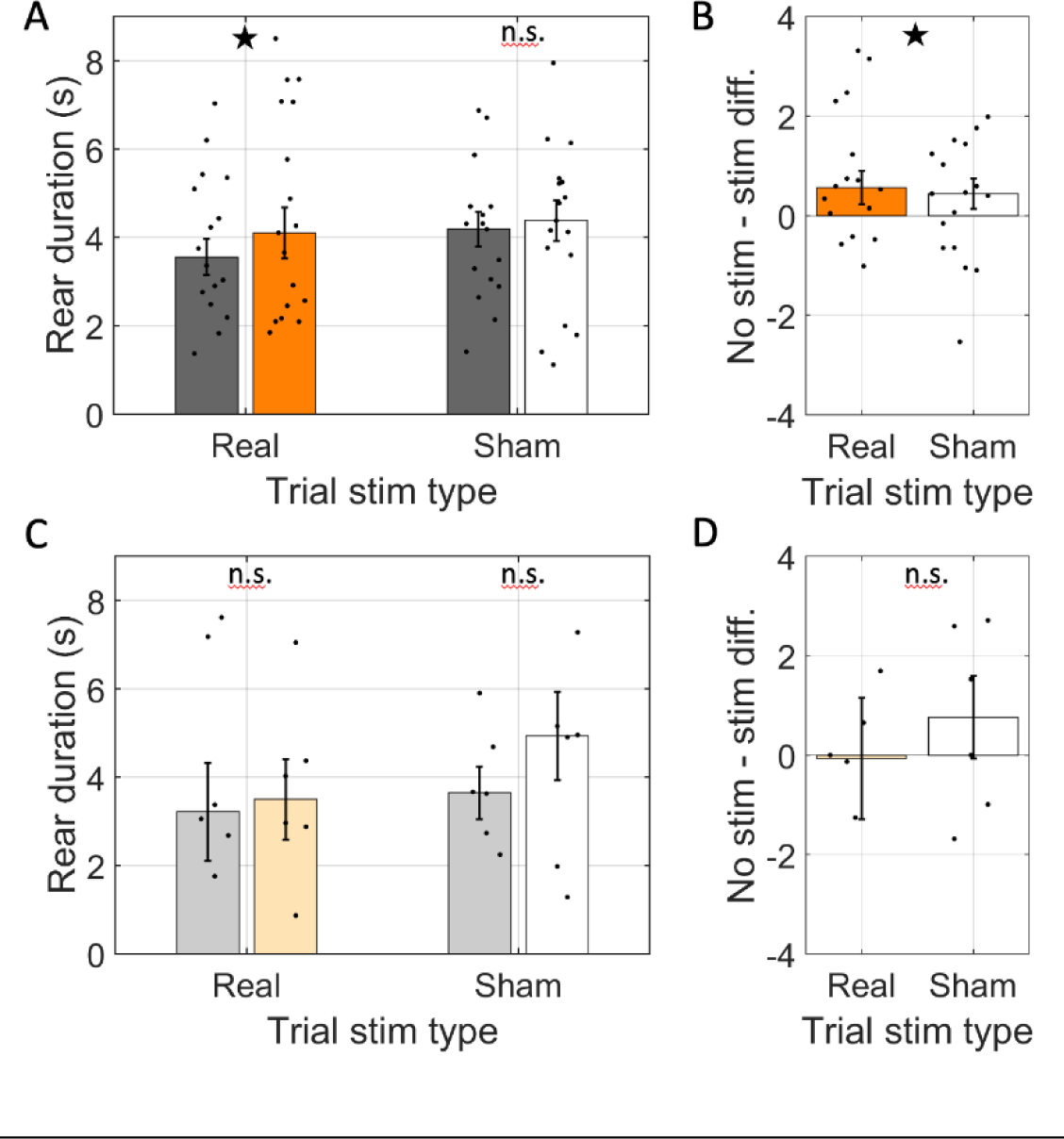
Rear duration was increased by optogenetic inhibition. A. The average duration of individual rears across rats transfected with halorhodopsin (n=17). Rear duration shown separately for rears in arms with simulation (real stimulation: orange, sham stimulation: white) and arms without stimulation (gray) on trials with the laser powered on (Trial stim type: real) and trials with the laser powered off (Trial stim type: sham). B. The difference in average rear duration between arms with and without stimulation in real and sham stimulation trials. Average rear duration was significantly greater in stim arms only in real stimulation trials. C & D. Same as A & B but for rats transfected with reporter only (n=6). In all panels, dots indicate the mean over trials for individual rats, bar height indicates the median over rats, and error bars indicate standard error over rats. ★ indicates p < 0.05. n.s. indicates *p* > 0.05.

The increase in rear duration in real stimulation arms was specific to the rats transfected with halorhodopsin. In rats transfected with reporter only, the duration of rears in real-stim-arms (glme est. = 3.9 s) was not significantly different from those in no-stim-arms (4.4 s; glme est. = - 0.49 [-2.53 1.56], *t*(50) = -0.48, *p* = 0.63; Fig. 3C & 3D). Likewise, sham stimulation did not affect rear duration (sham-stim = 4.7 s vs. no-stim = 3.9 s; glme est. = 0.79 [-1.13 2.71], *t*(51) *=* 0.83, *p* = 0.41; Fig. 3C & 3D) and no difference was found between real-stim and sham-stim trials (glme est. = 0.14 [-1.22 1.50], *t*(103) = 0.2, *p* = 0.84). The significant increase in rear duration during optogenetic inhibition of medial septal ChAT neurons indicates that cholinergic tone supports processing during rearing rather than generate the behavioral state that motivates rearing. In a hypo-cholinergic state, the rats take more time to complete the rear-related processing.

Though our experiment was motivated by the correlations between rearing and cholinergic tone, there are other behavioral markers worth investigating in the context of the present experiment as they offer further insights into the behavioral state of the animal. These include distance traversed in each arm and dwell times in each arm.

Distance traveled provides a window into whether the active locomotion of the rats was affected by the optogenetic manipulation of the medial septal ChAT neurons. Comparing the average cumulative distance traveled in real-stim and no-stim arms revealed that the cumulative distance traveled in real-stim-arms (glme est. = 84 cm) was not significantly different from the distance traveled in no-stim-arms (77 cm; glme est. = 6.6 [-1.1 14.4], *t*(261) = 1.68, *p* = 0.09; Fig. 4A & 4B). Likewise, distance traveled did not differ significantly in any other contrast.

**Figure 4:**
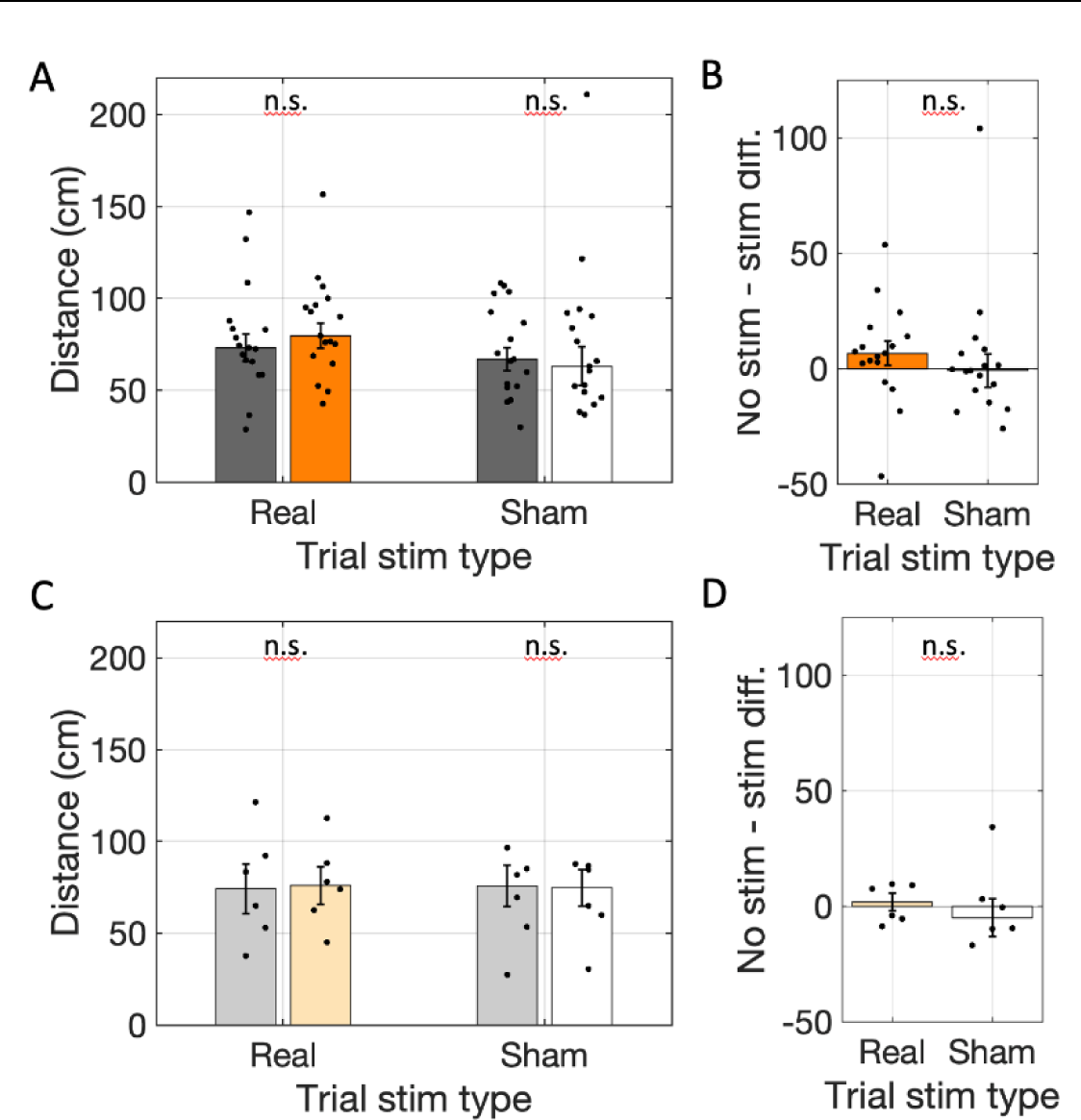
Total distance traveled did not change with optogenetic inhibition. A. The cumulative distance traveled in real-stim-arms (orange) was not significantly different from in no-stim-arms (gray) across rats transfected with halorhodopsin (n=17) in trials with the laser powered on (Trial stim type: real) or trials with the laser powered off (Trial stim type: sham). B. The differences in distance traveled between arms with and without stimulation in real and sham stimulation trials were not significantly different. C & D. Same as A & B but for rats transfected with reporter only (n=6). In all panels, dots indicate the mean over trials for individual rats, bar height indicates the median over rats, and error bars indicate standard error over rats. n.s. indicates *p* > 0.05.

Explicitly, the distance traveled in sham-stim-arms (77 cm) was not significantly different from no-stim-arms (72 cm; glme est. = 5.6 [-3.5 14.7], *t*(310) = 1.22, *p* = 0.22; Fig. 4A & 4B) and sham-stim trials were not significantly different from real-stim trials (glme est. = 5.9 [-0.62 12.5], *t*(573) = 1.78, *p* = 0.08). Further, the distance traveled in reporter-only rats did not differ between the real-stim trials (real-stim = 77 cm and no stim-stim = 75 cm; glme est. = 1.5 [-15.0 18.0], *t*(56) = 0.18, *p* = 0.86; Fig. 4C & 4D) nor in the sham-stim trials (sham-stim = 69 cm vs. no stim-stim = 69 cm; glme est. = 0.4 [-15.1 15.9], *t*(56) = 0.05, *p* = 0.96; Fig. 4C & 4D) and there was no difference between the trial types in the reporter-only cohort (glme est. = 0.96 [-10.6 12.5], *t*(114) = 0.16, *p* = 0.87). The lack of changes in distance traveled indicates that the inhibition of medial septal ChAT neurons did not affect the locomotor behavior of the rats.

To further test for changes in the behavioral state of the animals we asked if the total time rats spent in each arm (i.e., dwell time) changed. Dwell time, however, was not different between real-stim-arms and no-stim-arms (real-stim = 25.3 s; no stim = 22.6 s; glme est. = 2.69 [-1.46 6.84], *t*(261) = 1.28, *p* = 0.20; Fig. 5A & 5B). No differences were expected in the other conditions, and none were observed. Explicitly, there was no difference between arm types in sham-trials (sham-stim = 16.9; no-stim = 16.2; glme est. = 0.65 [-1.94 3.24], *t*(310) = 0.49, *p* = 0.62; Fig. 5A & 5B) nor did the reporter-only rats have differences in either real-stim trials (real-stim = 14.1; no-stim = 17.3; glme est. = -3.17 [-7.9 1.6], *t*(56) = -1.3, *p* = 0.19; Fig. 5C & 5D) or the sham-stim trials (sham-stim = 19.0; no-stim = 22.9; glme est. = -3.85 [-17.4 9.7], *t*(56) = 0.57, *p* = 0.57; Fig. 5C & 5D). The lack of difference in total dwell time indicates that the rats did not spend significantly longer in the arms.

**Figure 5:**
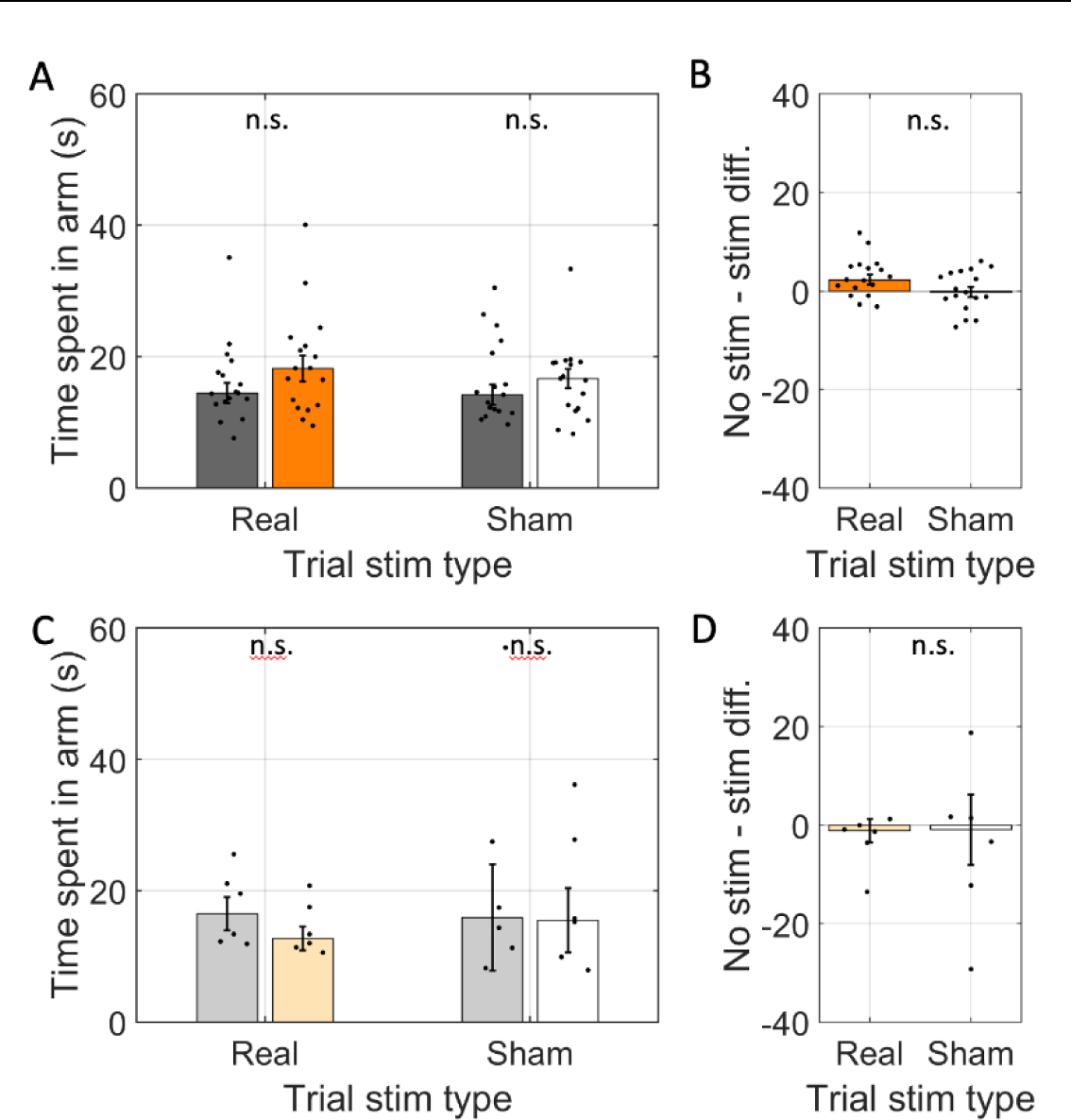
Average time spent in individual arms (i.e., dwell time) was not changed by optogenetic inhibition. A. The average time spent in individual arms across rats transfected with halorhodopsin (n=17). The time spent in individual arms is shown separately for arms with simulation (real stimulation: orange, sham stimulation: white) and arms without stimulation (gray) on trials with the laser powered on (Trial stim type: real) and trials with the laser powered off (Trial stim type: sham). B. The difference in time spent in arms with and without stimulation in real and sham stimulation trials. Average time spent was not significantly between arm types and was statistically indistinguishable between trial types. C & D. Same as A & B but for rats transfected with reporter only (n=6). In all panels, dots indicate the mean over trials for individual rats, bar height indicates the median over rats, and error bars indicate standard error over rats. n.s. indicates *p* > 0.05.

Finally, given acetylcholine’s role in encoding spatial memories and the assumed role of rearing in spatial orientation, we tested if modulating cholinergic neuron activity impacted task performance. To do so, we compared test phase performance between real-stim-trials and sham-stim-trials. We first looked at accuracy, scored as the percentage of first four arm entries were correct (rewarded). Accuracy on real-stim-trials (86%) was not significantly different from sham-stim trials (86%; glme est. = 0.8% [-2.6% 4.1%], *t*(305) = 0.45, *p* = 0.65; Fig. 6A). When rats miss one or more rewards in the first four entries it can take different amounts of time to locate the missed rewards. Quantifying the total number of arm entries to collect all four rewards captures this variability. However, total arm entries also did not differ (real-stim trials = 5.2 arms vs. sham-stim trials = 5.0 arms; glme est. = 0.26 arms [-0.21 0.74], *t*(305) = 1.09, *p* = 0.28; Fig. 6B). To test if the few errors made occurred on stim-arms disproportionately often, we separated the errors from real-stim trials based on the stim status at study. Again, we found no significant difference (0.54 errors were in stim-arms vs. 0.44 were in no-stim-arms; glme est. = 0.09 [-0.15 0.33], *t*(276) = 0.77, *p* = 0.44; Fig. 6C). No differences were expected across conditions in the reporter-only rats, none were found. Explicitly, accuracy was not statistically different (83% for real-stim vs. 86% for sham-stim; glme est. = -3.1 [-9.2 3.1], *t*(125) = -0.99, *p* = 0.32; Fig. 6D), total arm entries were not statistically different (5.2 arms for real-stim vs. 5.3 arms for sham-stim; glme est. = -0.07 arms [-0.82 0.67], *t*(125) = -0.20, *p* = 0.85; Fig 6E), and errors in real-stim trials were not statistically different between stim-arms and no-stim-arms (0.51 errors in stim-arms vs. 0.48 error in no-stim-arms; glme est. = 0.03 [-0.24 0.31], *t*(124) = 0.23, *p* = 0.82; Fig. 6E). These analyses of memory performance indicate that spatial memory performance was not impacted by the optogenetic manipulation.

**Figure 6:**
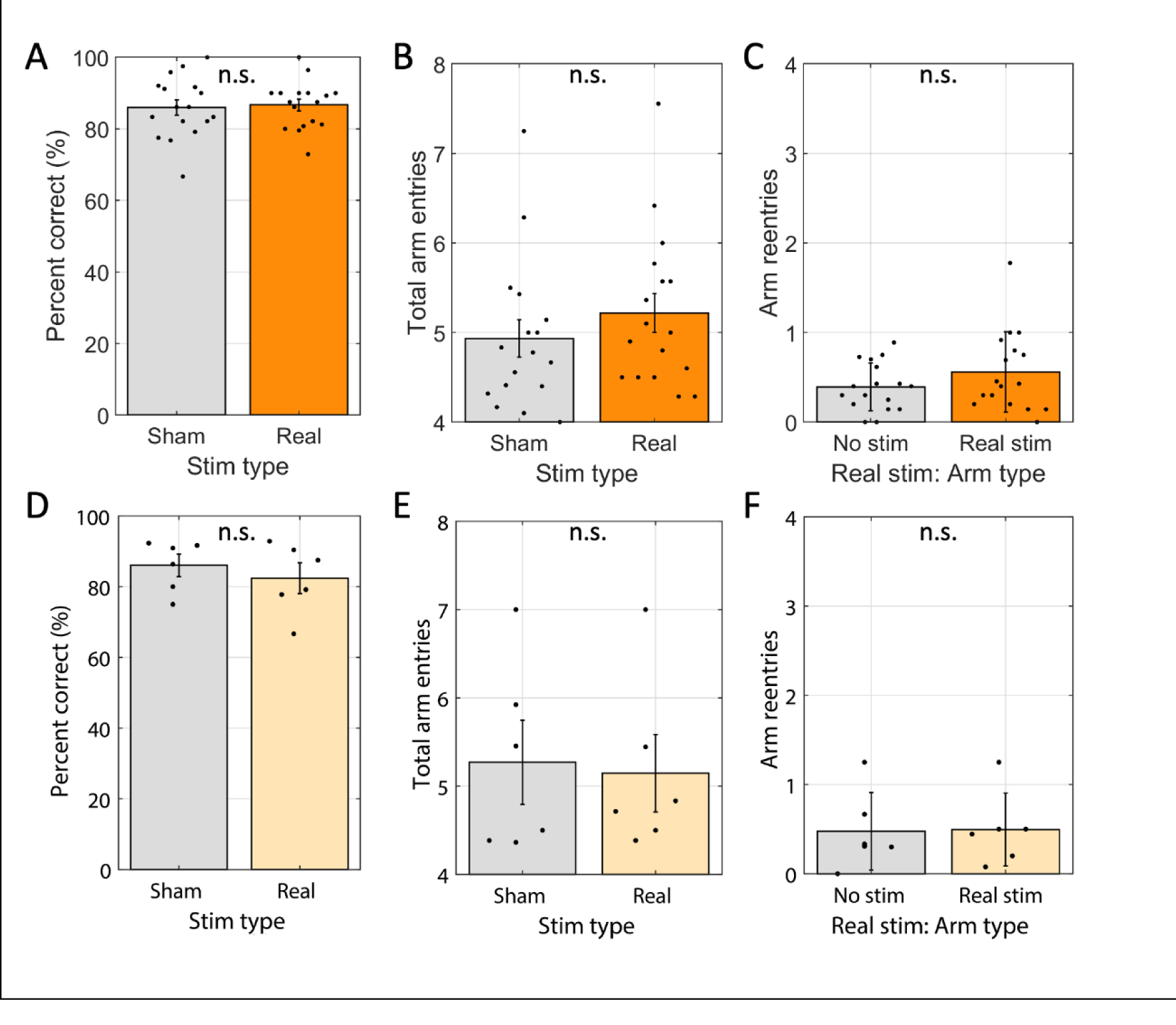
Win-shift performance was not changed by optogenetic inhibition. A. Average percentage of the first four arms entered containing reward (Percent correct) for sham stimulation trials (‘sham’) and real stimulation trials (‘real’) were statistically indistinguishable across rats transfected with halorhodopsin (n=17). B. Average number of arms entered to find the four rewards in sham stimulation and real stimulation trials were statistically indistinguishable across same rats as shown in A. C. For real stimulation trials, the average number of times each rat entered a no-stim arm relative to real stim arms were statistically indistinguishable across same rats shown in A. D–F. Same as A-C, respectively, but for rats that were transfected with reporter only (n = 6). Dots indicate the average over trials for individual rats, bar height indicates the mean over rats, and error bars indicate standard error over rats. n.s. indicates p > 0.05.

Given prior reports of context specific effects of manipulations of cholinergic signaling, we examined whether exploratory behavior differed in a context outside of the memory demands of the 8-arm win-shift task by activating the laser in the pre-trial hub-retention phase. As with real-stim and sham-stim trials, rats were held in the maze hub for up to two-minutes before any doors opened. Rats standardly rear frequently in this phase. ‘Hub-stim trials’ activated the laser throughout this retention phase and off-line analyses examined whether measures of exploratory behavior, as above, were affected.

As with modulation of cholinergic neuron activity in the arms, rearing frequency did not change (4.0 rears in hub-stim trials vs. 4.4 rears in sham-stim trials; glme est. = -0.37 [-1.09 0.35], *t*(193) = -1.01, *p* = 0.31; Fig 7A). For hub-stim trials, however, we found no change in rear duration (3.2 s in hub-stim trials vs. 3.4 s in sham-stim trials; glme est. = -0.18 [-0.90 0.53[, *t*(168) = -0.51, *p* = 0.61; Fig. 7B). We also found no changes in the distance traveled (266 cm in hub-stim trials vs. 264 cm in sham-stim trials; glme est. = 1.3 cm [-23.7 26.4], *t*(193) = 0.11, *p* = 0.92; Fig. 7C). Dwell time was in the hub was experimentally fixed and could not differ between conditions. No effect was expected among rats transfected with reporter only and none were found. Explicitly, rear count did not change (5.5 rears in hub-stim trials vs. 5.5 rears in sham-stim trials; glme est. = 0.03 rears [-1.6 1.7], *t*(54) = 0.04, *p* = 0.97; Fig. 7D), average rear duration did not change (2.2 s in hub-stim trials vs. 2.3 s in sham-stim trials; glme est. = -0.09 s [-0.65 0.46], *t*(48) = -0.33, *p* = 0.74; Fig 7E), and distance travelled did not change (135 cm in hub-stim trials vs. 140 cm in sham-stim trials; glme est. = -5.1 cm [-33.4 23.1], *t*(54) = 0.36, *p* = 0.71; Fig. 7E). These results indicate that there was no effect of inhibiting the cholinergic neurons of the medial septum on exploratory behaviors during the pre-trial phase.

**Figure 7:**
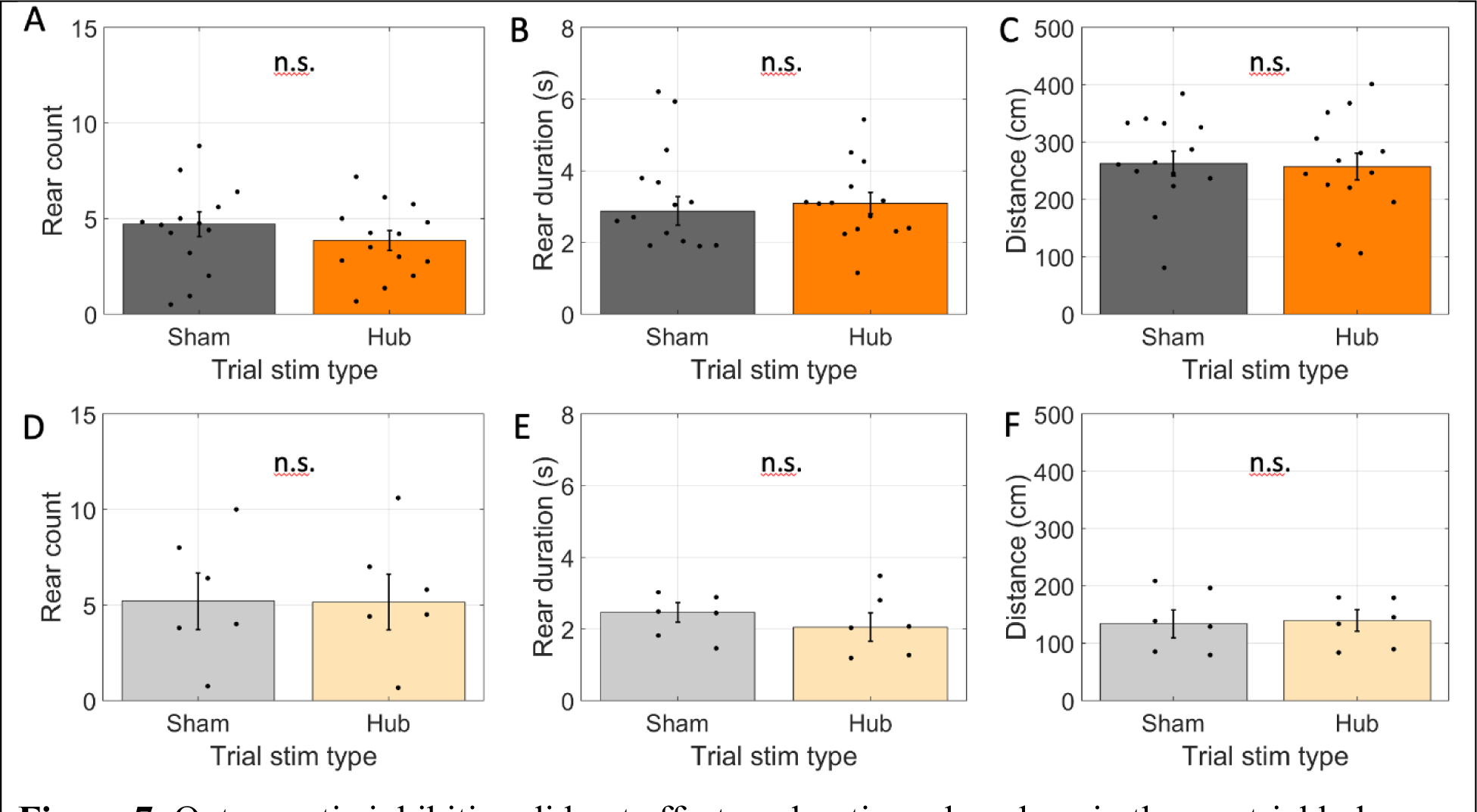
Optogenetic inhibition did not affect exploration when done in the pre-trial hub retention phase. A-C. Results for rats transfected with halorhodopsin. A. The number of rears during the pre-trial hub retention phase in trials with no laser stimulation (sham) and laser stimulation throughout the phase (Hub). B. The average duration of individual rearing events during the pre-trial hub retention phase. C. The total distance traveled during the pre-trial hub retention phase. D-F. Same as A-C but for rats transfected with reporter only. The average time spent in the maze hub was not significantly different between stim and sham trials. In all panels, dots indicate the average over trials for individual rats, bar height indicates the mean over rats, and error bars indicate standard error over rats. n.s. indicates p > 0.05.

## Discussion

This project aimed to test if the activity of cholinergic cells of the medial septum promoted rearing behavior in the context of a spatial working memory task. This hypothesis was motivated by previous correlational studies of exploration and cholinergic tone (Flicker & Geyer, 1982; van Abeelen, 1989; Acquas et al., 1996; Thiel et al., 1998; Giovannini et al., 2001; see Lever et al., 2006 for a related review). Such a connection, if it existed, could provide an explanation for the association between cholinergic modulation and spatial memory encoding given that rearing itself is a key epoch of spatial memory encoding (Layfield et al., 2023). Arguing against this connection however, we found that optogenetically inhibiting the activity of cholinergic neurons in the medial septum did not change rearing frequency. Rather, the optogenetic inhibition significantly increased rearing duration and total distance traveled.

The pattern of empirical findings reported here, a combination of no change in rearing frequency and an increase in rearing duration and movement in stimulation arms, argue against a simplistic hypothesis that activity of cholinergic neurons in the medial septum motivates an exploratory behavioral state. Such a hypothesis would have predicted reductions in both exploration and rearing.

Nonetheless, these results should not be interpreted as indicating that there is no relationship between acetylcholine and exploration. Indeed, given the rich literature on the importance of acetylcholine for encoding (for relevant reviews, see Hasselmo, 2006; Easton et al., 2012; Newman et al., 2012; Honey, Newman, Schapiro, 2017) and dynamics of the hippocampal formation (Venditto et al., 2019, Newman et al., 2017, Newman, Climer, Hasselmo, 2014, Newman et al., 2013), it is reasonable to hypothesize that the importance of acetylcholine is down-stream rather than up-stream of the exploration. That is, acetylcholine does not motivate information collection but rather supports the effective encoding of the information once collected. By this view, the observed increase in rear duration during optogenetic silencing of the cholinergic neurons reflects the increased time needed to encode the necessary information. By analogy, this is like an increased reaction time to a more demanding stimulus. It takes longer to reach the same end state. The fact that we observed no significant change in behavioral performance in stim trials is consistent with the idea that the rats indeed reached the same end state.

It is worth noting, however, that there has been some debate about the role of cholinergic input from the medial septum for hippocampal-dependent memory encoding (e.g., Parent & Baxter, 2004). Immunotoxic lesions that selectively kill acetylcholine producing neurons of the medial septum can fail to produce substantial encoding deficits. Our lack of behavioral performance decline aligns with this finding. Yet, given the significant increase in rear duration and distance traveled, we reasoned that the lack of change in performance was due to adaptive compensation by the rats. The same may be true in other cases where immunotoxic lesions of cholinergic neurons in the medial septum failed to generate behavioral changes. Such lesions, after all, leave most of the cholinergic modulation in the hippocampus intact (Chang & Gold, 2004). As such, it leaves open the possibility that other systems adaptively compensate for reduced cholinergic tone in the hippocampus.

The optogenetic inhibition used here should be regarded as a decrease in cholinergic tone, not a silencing of cholinergic tone. This is because the *ChAT:: CRE+* rats used to enable expression of halorhodopsin selectively in acetylcholine producing neurons have been found to only express CRE in ∼50% of the ChAT+ neurons of the medial septum (Witten et al., 2011). While complete silencing would also have been fine, the objective of this study was achievable using the graded decrease.

The current pattern of results may seem at odds with a previous study cholinergic modulation of rearing described by Monmaur et al. (1997). Monmaur et al. reported that rearing frequency increased following a local infusion of the cholinergic agonist carbachol into the septum in rats being held in a narrow chamber (i.e., without a task). However, there are several key procedural differences to the current work. First, Monmaur et al. infused carbachol into the medial septum whereas we optogenetically modulated the ChAT expressing neurons of the medial septum. Crucially, the primary effect of the carbachol infusion on cholinergic signaling would be local and any broader effects would be secondary to those on the activity of the medial septum. In contrast, by modulating the ChAT neurons, which include projection neurons targeting the hippocampus and entorhinal cortex, our manipulation would impact a broader circuit. Further, the extent of influence local to the medial septum may have been smaller in our work relative to that resulting from the carbachol infusion. The medial septum broadly expresses cholinergic receptors across local cell types (Van der Zee & Luiten, 1994) and, crucially, receives cholinergic input from not only the local cholinergic neurons but also ascending input from the pedunculopontine and laterodorsal nucleus (Jones & Cuello, 1989). Thus, the optogenetic manipulations would only be expected to target some of the local targets. Another major difference between the findings of Monmaur et al. and those reported here is our inclusion of a task. Thus, it is unclear what behavioral motivation the rearing Monmaur et al. observed may have been related to. For example, the rats may have been focused on looking for opportunities to escape. Here, we were specifically interested in spatial memory encoding.

Comparing the present results to those of Thiel et al., 1998 also provides a useful counterpoint. In brief, Thiel et al. used microdialysis to examine acetylcholine release in rats as they habituated to an open field and compared the release to the extent of exploratory behavior. They found increased ACh release while the rat was in the open field. On the first exposure to the open field, when compared over rats, rearing frequency was positively correlated with acetylcholine release. On re-exposure to the open field, rearing was unrelated to acetylcholine release. In the present work, we found no change in rearing frequency when we modulated the activity of the cholinergic neurons. This aligns best with what Thiel et al found in rats being re-exposed to an enclosure in that our rats, which were habituated to the testing enclosure, also had no relationship between rearing frequency and cholinergic modulation. Beyond examining rearing frequency, Thiel et al. also examined locomotion (virtual line crossings). They found no correlation with acetylcholine release. This is at odds with our finding that rats traveled further in arms where the cholinergic neurons were inhibited. However, as in the study by Monmaur et al., the animals examined by Theil et al. were not performing a task. The varying behavioral context (e.g., presence versus absence of task demands) thus appears to modulate the relationship between cholinergic tone and exploration.

Supporting the idea that differences in behavioral context could impact the relationship between cholinergic tone and rearing, we found here that optogenetic inhibition only increased rear duration and distance traveled in when rats were underway in the study phase. When we repeated the optogenetic inhibition manipulation while the rats waited for the study phase to start in the maze hub, we found no change in either rear duration or distance traveled. By this line of thinking, rearing in the maze hub prior to the start of the study phase may be motivated by a different behavioral motivation than rearing that occurs in the arms once the study phase has started. Prior to the trial start, because all arms are baited and the rat cannot know which doors will open, rearing provides minimal diagnostic information about future actions. However, once the trial has started and the rat is in an arm, rearing can offer useful trial specific information about where it has been or where it may be going. Given this difference in the utility of rearing, it is conceivable that rearing is driven by different motivations in the different contexts with differing physiological mechanisms.

In summary, the present findings show that inhibiting these neurons has no effect on rearing frequency but increases rearing duration. This argues against the hypothesis that the activity of acetylcholine producing neurons in the medial septum motivates an exploratory behavioral state. Rather, these results suggest that acetylcholine is functionally related to exploratory behavior because it is triggered by common upstream mechanisms and supports the processing of information acquired during exploration. As such, the observed increase in duration could be explained by the rats requiring more time to perform the same encoding. Finally, the current results support the hypothesis that rearing is multiply determined as the same optogenetic inhibition manipulation had different effects on rearing in different behavioral contexts.

## Acknowledgements

We would like to thank from Drs., Jonathon Crystal and Preston Garraghty for input and feedback on the project and manuscript. This research was supported in part by Lilly Endowment, Inc., through its support for the Indiana University Pervasive Technology Institute. We are grateful to the care and attention the IU Laboratory Animal Resources (LAR) core facility give every day to the rats at the heart of this project. This project was supported by Bridge Funds from the IU OVPR and NIH R01 AG076198. SC was supported by The Brotheridge Family IU, Office of Undergraduate Research and IU Hutton Honors College. IC was supported by the Brotheridge Family IU and IU Hutton Honors College. SB and LO were supported by IU Hutton Honors College. AI and KM were supported by IN LSAMP. DL was supported by the Harlan Scholars Program and the IU PBS Neuroscience Fund.

## Abbreviations

ChAT: Choline acetyltransferase
EYFP: Enhanced Yellow Fluorescent Protein
GFP: Green Fluorescent Protein
PBS: Phosphate Buffered Saline
n.s.: Not significant (*p* > 0.05)
glme: generalized linear mixed effects

## Author contributions

The authors confirm contribution to the paper as follows: SC, DL, ELN conceived and designed the experiment methodology. SC, BHG, AMI, SCB, KTM, AES, LO performed the investigation. SC, IJC, ELN conceived of and implemented the formal analyses, performed the visualization and wrote the original manuscript. IJC and ELN performed the writing review and editing.

## References

1. Abeelen, J. H. F. van. (1989). Genetic control of hippocampal cholinergic and dynorphinergic mechanisms regulating novelty-induced exploratory behavior in house mice. Experientia 45, 839–845.

2. Acquas, E., Wilson, C. & Fibiger, H. C. (1996). Conditioned and Unconditioned Stimuli Increase Frontal Cortical and Hippocampal Acetylcholine Release: Effects of Novelty, Habituation, and Fear. J Neurosci 16, 3089–3096.

3. Blokland, A., Honig, W. & Raaijmakers, W.G.M. (1992). Effects of intra-hippocampal scopolamine injections in a repeated spatial acquisition task in the rat. Psychopharmacology 109, 373–376.

4. Dannenberg H, Pabst, M., Braganza O, Schoch S, Niediek J. (2015). Synergy of direct and indirect cholinergic septo-hippocampal pathways coordinates firing in hippocampal networks. The Journal of neuroscience : the official journal of the Society for Neuroscience 35, 8394–8410.

5. Douchamps, V., Jeewajee, A., Blundell, P., Burgess, N. & Lever, C. (2013). Evidence for Encoding versus Retrieval Scheduling in the Hippocampus by Theta Phase and Acetylcholine. The Journal of neuroscience : the official journal of the Society for Neuroscience 33, 8689–8704.

6. Easton, A., Douchamps, V., Eacott, M. & Lever, C. (2012). A specific role for septohippocampal acetylcholine in memory? Neuropsychologia 50, 3156–3168.

7. Elvander, E., Schött, P. A., Sandin, J., Bjelke, B., Kehr, J., Yoshitake, T., & Ögren, S. O. (2004). Intraseptal muscarinic ligands and galanin: influence on hippocampal acetylcholine and cognition. Neuroscience, 126(3), 541–557.

8. Flicker, C., & Geyer, M. A. (1982). Behavior during hippocampal microinfusions. II. Muscarinic locomotor activation. Brain Research Reviews, 4(1), 105–127.

9. Giocomo, L. M. & Hasselmo, M. E. (2005) Nicotinic modulation of glutamatergic synaptic transmission in region CA3 of the hippocampus. Eur J Neurosci 22, 1349–1356.

10. Gradinaru V, Zhang F, Ramakrishnan C, Mattis J, Prakash R, Diester I, Goshen I, Thompson KR, Deisseroth K. (2010) Molecular and cellular approaches for diversifying and extending optogenetics. Cell, 141(1):154–65.

11. Giovannini MG, Rakovska A, Benton RS, Pazzagli M, Bianchi, L, Pepeu G. (2001). Effects of novelty and habituation on acetylcholine, GABA, and glutamate release from the frontal cortex and hippocampus of freely moving rats. Neuroscience 106, 43–53.

12. Hasselmo M. E. (2006). The role of acetylcholine in learning and memory. Current Opinion in Neurobiology, 16(6), 710–715.

13. Hasselmo, M. E. & Schnell, E. (1994). Laminar selectivity of the cholinergic suppression of synaptic transmission in rat hippocampal region CA1: computational modeling and brain slice physiology. The Journal of Neuroscience 14, 3898–3914.

14. Honey, C. J., Newman, E. L. & Schapiro, A. C. (2017) Switching between internal and external modes: A multiscale learning principle. Network Neuroscience 1, 339–356.

15. Jones, B. E., and Cuello, A. C. (1989). Afferents to the basal forebrain cholinergic cell area from pontomesencephalic catecholamine, serotonin, and acetylcholine neurons. Neuroscience 31, 37–61.

16. Layfield D, Sidell N, Abdullahi A, Newman EL. (2020). Dorsal hippocampus not always necessary in a radial arm maze delayed win-shift task. Hippocampus 30, 121–129.

17. Layfield, D., Sidell, N., Blankenberger, K. & Newman, E. L. (2023) Hippocampal inactivation during rearing on hind legs impairs spatial memory. Scientific Reports 13, 6136.

18. Lever, C., Burton, S., & O’Keefe, J. (2006). Rearing on hind legs, environmental novelty, and the hippocampal formation. Reviews in the Neurosciences, 17(1-2), 111–133.

19. Frotscher, M., & Léránth, C. (1985). Cholinergic innervation of the rat hippocampus as revealed by choline acetyltransferase immunocytochemistry: a combined light and electron microscopic study. The Journal of comparative neurology, 239(2), 237–246.

20. Martyn, A. C. et al. (2012) Elimination of the vesicular acetylcholine transporter in the forebrain causes hyperactivity and deficits in spatial memory and long-term potentiation. Proc National Acad Sci 109, 17651–17656.

21. Mathis, A., Mamidanna, P., Cury, K.M. et al. DeepLabCut: markerless pose estimation of user-defined body parts with deep learning. Nat Neurosci 21, 1281–1289 (2018).

22. Monmaur P, Sharif A, M’Harzi M. (1997). Involvement of septal muscarinic receptors in cholinergically mediated changes in rat rearing activity. Pharmacology Biochemistry and Behavior, 58, 577–582.

23. Nath, T. et al. (2019) Using DeepLabCut for 3D markerless pose estimation across species and behaviors. Nat Protoc 14, 2152–2176.

24. Newman EL, Climer JR, Hasselmo ME. (2014) Grid cell spatial tuning reduced following systemic muscarinic receptor blockade. Hippocampus 24(6):643–55.

25. Newman EL, Gillet SN, Climer JR, Hasselmo ME (2013) Cholinergic Blockade Reduces Theta-Gamma Phase Amplitude Coupling and Speed Modulation of Theta Frequency Consistent with Behavioral Effects on Encoding. The Journal of neuroscience : the official journal of the Society for Neuroscience 33, 19635–19646.

26. Newman EL, Gupta K, Climer JR, Monaghan CK, Hasselmo ME. (2012) Cholinergic modulation of cognitive processing: insights drawn from computational models. Frontiers in behavioral neuroscience 6, 24.

27. Newman EL, Venditto SJC, Climer JR, Petter EA, Gillet SN, Levy S. (2017) Precise spike timing dynamics of hippocampal place cell activity sensitive to cholinergic disruption. Hippocampus 27(10):1069–1082

28. Olton, D. S. & Samuelson, R. J. (1976). Remembrance of Places Passed - Spatial Memory in Rats. Journal of Experimental Psychology-Animal Behavior Processes 2, 97–116.

29. Parent, M. B. & Baxter, M. G. (2004) Septohippocampal Acetylcholine: Involved in but not Necessary for Learning and Memory? Learn Memory 11, 9–20.

30. Thiel, C. M., Huston, J. P. & Schwarting, R. K. W. (1998). Hippocampal acetylcholine and habituation learning. Neuroscience 85, 1253–1262.

31. Venditto SJC, Le B, Newman EL. (2019) Place cell assemblies remain intact, despite reduced phase precession, after cholinergic disruption. Hippocampus 29(11):1075–1090.

32. Whishaw, I. Q., O’Connor, W. T. & Dunnett, S. B. (1985) Disruption of central cholinergic systems in the rat by basal forebrain lesions or atropine: effects on feeding, sensorimotor behaviour, locomotor activity and spatial navigation. Behavioural brain research 17, 103–115.

33. Witten, I. B., Steinberg, E. E., Lee, S. Y., Davidson, T. J., Zalocusky, K. A., Brodsky, M., Yizhar, O., Cho, S. L., Gong, S., Ramakrishnan, C., Stuber, G. D., Tye, K. M., Janak, P. H., Deisseroth, K. (2011) . Recombinase - driver rat lines: tools, techniques, and optogenetic application to dopamine - mediated reinforcement. 72, 721–73.

